# *Social clues in risky pools*: Perceived conspecific rivals strongly modify individual oviposition decisions in response to larval predation and competition

**DOI:** 10.1101/2024.07.03.602002

**Authors:** Ashwini Ramesh, Manvi Sharma, Kavita Isvaran

**Affiliations:** Indian Institute of Science, Bengaluru, India – 560071; Indian Institute of Science Education and Research, Pune, India – 411008; Michigan State University, East Lansing, USA – 48823; Ashoka University, Sonipat, India – 560100

**Keywords:** social information, social facilitation, social inhibition, oviposition site selection, predation, competition, mosquito, *Aedes aegypti*

## Abstract

Reproductive choices are imperative in shaping organismal fitness across diverse taxa. Such choices are particularly critical in organisms with biphasic lifecycles, as females must maximize offspring survival pre-oviposition, with no parental care extended afterward. Consequently, females face strong site selection pressures to effectively respond to offspring competition and predation risks. Ovipositing females encounter yet another challenge during site selection: assessing future competition for their offspring from potential conspecific rivals. Our current knowledge, based on average social versus solitary behaviours, fails to clarify how social signals influence individual behavior within groups. To address these challenges, we leveraged the unique oviposition biology of the mosquito *Aedes aegypti* where only blood-fed females can lay eggs. By tracking individual behaviour in a social setting, we ask: how does social information from perceived conspecific rivals influence an individual’s oviposition site selection? In our lab-based experiment, we examined oviposition strategies at two spatial scales under varying larval competition and predation risk. Our findings reveal that social information exerts a stronger influence on egg-laying behavior at larger spatial scales, i.e., at the scale of pool networks, than between neighboring pools. Social cues facilitated oviposition with increasing larval predation, as social females transitioned from rejecting to accepting pool networks. Conversely, under larval competition, social cues led to inhibition, with females withholding their eggs likely in anticipation of future competition. At finer spatial scales, social information only weakly modified oviposition behavior despite potential negative fitness consequences for the offspring. Thus, perceived conspecific risk strongly modifies oviposition—facilitation, inhibition, or no effect—and is scale-dependent.

## INTRODUCTION

Reproductive site selection is arguably one of the strongest forces shaping organismal fitness in diverse taxa. Oviposition/nest site selection (OSS), a type of reproductive site selection, is the act of choosing a habitat for egg-laying (*reviewed in* Refsnider and Janzen 2010). Many ovipositing species typically deposit eggs or juveniles at sites unaccompanied by parental care (e.g. amphibians, mosquitos, nesting reptiles). Offspring at this stage are vulnerable to both competition and predation risk that has consequences for their survival, longevity, and reproduction (Mukherjee and Blaustein 2019; Gimnig et al 2002). For example, *A. aegypti* survivors from predator pools had shorter development times, larger wing spans, and significantly fewer offspring with increasing predator density (Sharma et al. 2020). Thus, females are expected to make choices that maximize offspring fitness prior to oviposition, as following this act no parental care can be extended to the offspring (Refsnider and Janzen 2010). Consequently, females face strong selection pressures to respond effectively to offspring competition and predation risk.

However, ovipositing females are confronted with an additional challenge: grappling with the implications of conspecific presence at oviposition sites. Social cues derived from conspecific behavior can significantly reshape both the process and outcomes of oviposition (*reviewed in* Buxton et al 2020). While the presence of conspecifics may signal a high-quality oviposition site, their potential oviposition may pose a competition threat to a female’s future offspring (Bonnie and Earley 2007; Betts et al 2008). The fluctuating densities of adult conspecifics across seasons dictate the social information available to ovipositing individuals (Miller et al 2013). In the early season, lower competition among breeding females may lead to solitary individuals at oviposition sites, where they encounter fewer larval competitors. Conversely, peak season witnesses increased interactions among ovipositing females, heightening larval competitors and their predators (Lehmann et al 2014). Consequently, individuals must simultaneously weigh both the potential competition from the offspring of perceived conspecific rivals and the immediate risks to their current larvae from other larvae or predators, which collectively can either hurt or harm their offspring (Drugowitsch et al. 2012; *reviewed in* Buxton et al. 2020). On one hand, while average oviposition behavior under solitary versus grouped conditions is well established (Prokopy and Bush 1973; Le Goff et al. 2010), the influence of social information on individual oviposition behavior within a group remains elusive. On the other hand, although individual responses to larval predation and competition risks are well studied, how social information modifies those risks is poorly understood. In non-social insects, where inclusive fitness benefits due to genetic relatedness are limited, investigating inter-individual variation in oviposition can yield crucial insights into individual fitness and population dynamics (Tóth et al 2020).

Unfortunately, the consequences of social information on individual oviposition decisions remain difficult to predict for several reasons. First, the effect of social information on oviposition via perceived conspecific rivals has produced inconsistent findings. In non-social insects, there are reports of females in a social setting showing increased egg-laying (social facilitation), or decreased egg-laying (social inhibition), or similar when compared to solitary females (Prokopy and Bush 1973; Chess et al 1990; Abernathy et al 1994; Le Goff et al 2010; Visser 1994; Aikins et al 2023). Second, constraints in study design have limited our ability to test inter-individual variation in oviposition response to social cues. Current understanding primarily relies on the average behavior of adults in social versus solitary settings, owing to challenges of monitoring the fecundity of individuals. Individuals are known to exhibit high intra- and inter-individual variation in egg-laying(Clements 1992; Gibbs et al 2005). Thus, in the absence of tracking known individuals, the reproductive success of social *v* solitary females remains elusive. Third, oviposition decisions are further compounded as females may respond differently to larval predation and competition risks. When faced with larval competition, females may avoid ovipositing in pools with conspecific broods to ensure more resources for their offspring (Munga et al. 2006). In contrast, under larval predation, there is evidence of both avoidance (Blaustein et al. 2004; Resetarits and Wilbur 1989) and preference for pools with predators across taxa (Sharma and Isvaran, 2019). Consequently, key life-history trade-offs may shift along these distinct selection gradients affecting fitness traits in both adults and offspring. Finally, the spatial scale at which social cues influence oviposition decisions remains elusive. Specifically, do social cues operate at finer spatial scales, modifying female egg-laying choices between neighbouring risky and risk-free sites? Or do social cues operate at larger scales, prompting females to reject a set of oviposition sites despite the availability of risk-free options at finer scales? If yes, then how do they operate across scales?

Here, we aim to resolve these challenges by studying the influence of social cues on an individual female’s oviposition response to pools varying in risk using the mosquito, *Aedes aegypti,* as a model system. Specifically, we test how social information from perceived conspecific rivals influenced an individual’s pattern of oviposition in response to pools varying in either larval competition or larval predation risk. By leveraging the unique biology of Aedes mosquitoes, where only blood-fed females can lay eggs, we can safely track and compare the oviposition behavior of individual females in social settings to those in solitary conditions. In a series of binary choice experiments, adult gravid *A. aegypti* mosquitoes were housed either individually (solitary) or in the presence of non-egg-laying adult females (social). They were presented with two pools: a risk-free control pool and a potentially risky treatment pool. We investigated oviposition responses at two scales: at the larger spatial scale, the control and treatment pools together represent a pool network; at the finer spatial scale, choices were made between the neighbouring risky and risk-free pools. The treatment pool contained either one of two sources of risk: larval competition and predation risk, each examined at two levels: low and high. A set of control and treatment pools is referred to with respect to the level and type of risk (e.g., a pool network consisting of a control pool and a low predator treatment pool is referred to as “low” background predation risk). We assumed that individual females perceived the presence of other females in their environment as potential competitors for egg-laying sites, thus increasing the perceived intensity of larval competition. In light of this ecology, we hypothesize that:

### A) Larval Competition

#### Large spatial scale

Under ‘*low*’ background larval competition, social females were expected to reject more oviposition sites and withhold more of their fecundity when they perceive adult ovipositing competitors compared to solitary females. This response is anticipated to be even stronger under ‘*high’* background larval competition. As females lay multiple clutches, withholding eggs enables them to exercise future egg-laying opportunities.

#### Fine spatial scale

Under ‘*low*’ background larval competition, social females should aim to minimize competition from current larval competitors in the treatment pool. Importantly, social females should seek to avoid additional competition arising from perceived conspecific rivals in the control pool. Thus, social females are expected to distribute their eggs across the pools, laying slightly more eggs in the control pool. Solitary females, in contrast and without the added risk from perceived conspecific rivals, are expected to lay more eggs in treatment pools, as it cues stable habitats (conspecific cueing). However, as background larval competition intensifies, this response is anticipated to be even stronger, resulting in a greater skew to oviposit in control pools in both social and solitary females.

### B) Larval Predation Risk

#### Large spatial scale

Under both ‘*low’* and ‘*high’* background larval predation risk, social females should accept more pool-networks and increase total fecundity compared with solitary females, with this effect becoming stronger as background predation intensifies. For an individual in a social setting, the per capita risk for her own offspring is likely to be low due to the presence of additional unrelated larvae contributed by other females in the environment, creating a potential dilution effect.

#### Fine spatial scale

Under low larval predation risk, social females may distribute their eggs between pools if the perceived dilution effect is not strong enough. However, under high predation risk, social females are expected to distribute their eggs less frequently, concentrating more eggs in predator pools due to the potential dilution effect. In comparison, solitary females are expected to consistently lay eggs in control pools, as the per capita risk for her offspring in any predator pool is likely to be higher than in a social setting

## METHODS

### Model System

In this study, we used a lab-bred colony of *A. aegypti* mosquitoes. *A. aegypti* undergoes complete metamorphosis with aquatic egg, larval, and pupal stages. Its relatively short life-span and even shorter breeding period rendered it easy to evaluate reproductive success and female behavior. Females of this species have been recorded to distribute eggs spatially and temporally and engage in super-oviposition behavior (i.e. ovipositing on the same surface where other conspecific females have deposited eggs). In addition, *Bradinopyga* nymphs, a common predator of *A. aegypti* larvae, were also easily accessible from tanks maintained in the nurseries on IISc campus. These dragon fly nymphs, whose aquatic phase lasts for about 30 days before they metamorphose as terrestrial adults, inhabit and oviposit in the same set of natural pools as do several species of mosquitoes. These predators are generally benthic or found on walls of the pools and found to heavily predate upon larvae of mosquitoes (Clements 1992).

### Study Design

To capture the influence of social information on oviposition decisions in the context of seasonal distribution of adult densities, we measured individual oviposition decisions in varied adult conspecific densities. In a laboratory setting, we created two assay environments – solitary and social. Individual adult females with blood fed distended abdomens were isolated from a caged colony of ~150 adult mosquitos. The individual females were isolated using aspirators and placed in individual cages (0.3 m x 0.3m x 0.3 m) that were randomly pre-assigned to either solitary or social status. In the solitary cages, a single blood fed female was caged alone while in the social cages, a single blood fed female was caged with three other non-blood fed females. Since non-blood fed females typically do not invest in egg-laying behavior, one could safely track the oviposition behavior of an individual female in a social setting and compare her behavior to that of an individual female in the solitary condition. Each of the assay cages was supplied with ad libitum food source, i.e., moist cotton smeared with honey, which was replenished regularly, and maintained at a 14:10 hour day and night cycle, at 27 ± 5 °C. On the third day from the introduction of the females into the assay cages, each cage was provided with two “ovicups” (4 cm x 11 cm) containing treatment and control cue-water.

For the treatment, we examined two sources of risks, larval predation and larval competition. Previous work in the lab examined larval growth and performance when different densities of larvae were reared together (competition) and when larvae were preyed upon by varying densities of predators (predation) (Sharma et al 2020). Based on the results from this study, we chose larval and predator densities that would represent low and high risk of competition and predation, respectively, for offspring of ovipositing mosquitoes. In the predation treatment, the low and high corresponded to 2 and 4 predators, and in the competition treatment 20 and 155 larvae. Three days prior to the experiment, porous bags containing 0.2g of fish food were soaked in 1.5 L tubs of water. For predator treatments, we simultaneously added predators to the tub. For competition treatments, we added the required number of 1^st^ instar larvae (reared separately) to the tub. No additions were made to the control tubs, which still contained the aforementioned 0.2g of fish food. One day prior to the experiment, the porous bags were removed from the tubs. Water lost due to evaporation or when drawn for the experiment was replaced. Predators were removed from the tub, fed 3-5 larvae every other day and placed back in the tub. Larvae that pupated during the experiment were replaced. On the third day from the introduction of the females into the assay cages, 100 ml of water drawn from these tubs (cue-water) were transferred to “ovi-cups”. The cups, each lined with ovistrips, were always placed at diagonally opposite ends of the cage and randomly assigned to one of the four possible combinations of positions. The same position was maintained within a cage throughout the trial. The ovistrips and cue-water were replenished on a daily-basis for a period of three days. Finally, the individuals from the trial were aspirated and stored in Eppendorf tubes. Assay cages were donated to the colony and the cue-water tubs were disassembled at the end of a block of trials.

A total of 206 trials were conducted across 22 blocks in solitary and social assay environments. Each block lasted for seven days and included preparation of cue-water for control and treatment conditions, maintenance of assay cages and the running of 2 to 4 trials of different control-treatment binary choice experiments. The different control-treatment choice experiments used in this study were interspersed in time to avoid any confounding effects of time on oviposition responses.

Ethical Note: The work we report conforms to the requirements of the Institute Animal Ethics Committee of the Indian Institute of Science, Bengaluru, where the work was carried out, and to the laws of India.

## ANALYSIS

The primary measure of oviposition behaviour was the number of eggs laid by an individual female. Ovistrip eggs from control and treatment pools were manually counted and recorded, at intervals of 24 hours over a period of three days. The eggs from all three days were summed up for each pool.

### Large Spatial Scale

To understand how adult females assessed pool-network in its entirety, we measured:

#### 1. Propensity to oviposit

Here we recorded whether a female laid eggs during the trial. Females, after possibly assessing both pool type (control and treatment) and social status, can choose to either oviposit or entirely avoid the set of pools. Thus, propensity to oviposit was used as a measure of the degree of choosiness at the larger spatial scale of the set of pools.

#### 2. Total fecundity

The total number of eggs laid by a female during a trial, calculated as the sum of eggs deposited in control and treatment pools. Total fecundity can also be a measure of choosiness at the level of pool-network, if females can successfully manipulate the total number of eggs laid during the trial after assessment of the two pools and social status.

### Finer spatial scale

To understand how eggs are distributed between pools and estimate choosiness among females, we calculated an oviposition activity index.

#### 1. Oviposition Activity Index (OAI)

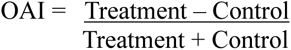

This index is a measure of the degree of choosiness shown by the female at the individual pool level. It identifies how eggs are distributed among pools (here between the control and treatment pools). This index, ranges from −1 to 1, where 1 indicates maximum preference for treatment (all eggs deposited in treatment pool alone), −1 maximum preference for control and 0 no preference (equal distribution of eggs across both pools). The absolute value of OAI indicates how choosy a female is, where values close to 0 indicate that females are more likely to distribute across pools, while those close to 1 indicate that females are highly choosy.

#### 2. Relationship between OAI and total fecundity

This relationship between OAI and total fecundity provides a unique insight into the relationship between the two spatial scales. Here we ask, how does the degree of egg distribution across pools impact a female’s decision of total number of eggs laid (total fecundity) in the pool-network?

To test the influence of social status on the oviposition responses of females to pools varying in risk, we ran separate linear mixed-effects models assuming relevant errors with each of the response measures. For propensity to oviposit, we ran generalized linear mixed models assuming binomial error with propensity to oviposit (oviposited or not) as the response variable. Risk level (low/high) and status (social/solitary) were modelled as fixed effects, and block (week of trial) as random effects. For each response variable, separate linear mixed-effect models were run for predation and competition risk. In this context, a set of control and treatment pools is referred to with respect to the level and type of risk (e.g., a pool network consisting of a control pool and a low predator treatment pool is referred to as “low” background predation risk). For total fecundity and OAI we ran separate linear mixed-effects models, assuming negative binomial and gaussian errors, respectively.. Risk level, and status were modelled as fixed effects, and block as random effects. Separate models were run for each source of risk.

To examine the relationship between OAI and total fecundity, we ran a generalized linear model in the family gaussian with total fecundity as the response variable and oviposition activity index (OAI), risk level, source of risk, status and the interaction term between OAI and status as the explanatory variables. We also included the quadratic term of OAI and the interaction term of the quadratic variable and status in the model. We included the interaction term in the model because we expected female response to vary with social status depending on the value of OAI.

For each analysis, cautious model reduction was performed. Only non-significant (p-value > 0.05) interaction terms were removed from the maximum model to aid in better understanding of relationships. Thus, inferences were based on a final model inclusive of all main effects (statistically significant or not), random effects, and interaction terms that were statistically significant. Additionally, to support our inferences about the strength of a relationship, effect sizes and bootstrapped confidence intervals were calculated.

The R version 4.4.0 (R Core Team, 2021) was used for all analyses and lme4 package (Bates et al 2015) to run mixed-effects models.

## RESULTS

### Larval Competition

#### Larger spatial scale

Overall, social information did not influence the propensity to oviposit (Fig. 1A, risk level: *X^2^* = 0.02983, *p* = 0.8629; status: *X^2^* = 0.43119, *p* = 0.5114) but strongly influenced total fecundity of social females across varying background larval competition (Fig. 1B, risk level: X^2^ = 0.2028, p = 0.6525; status: X^2^ = 3.14171, p = 0.002*). In both *‘Low’* and *‘High’* background competition, social females were as likely to oviposit in the pool network as solitary females but withheld their eggs. In *‘Low’* competition, they laid an average of 30 ± 2.3 eggs, significantly fewer than solitary females. As background larval competition intensified, social females laid an average of 36 ± 3.4 eggs, representing 0.89 times fewer eggs than those laid by solitary females.

**Figure 1:**
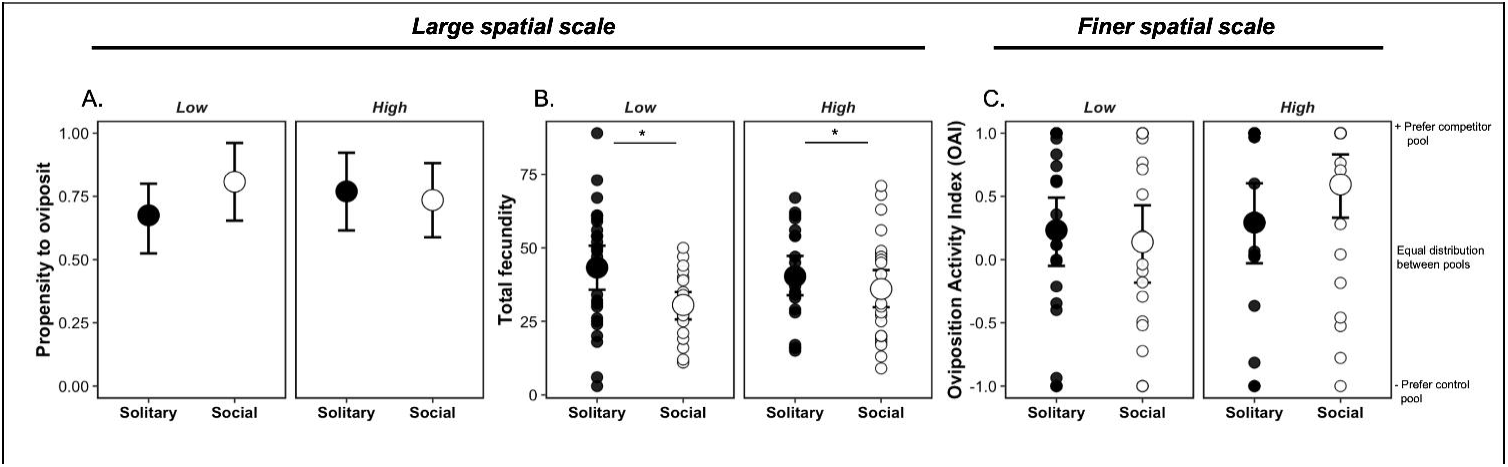
In both ‘*Low’* and ‘*High*’ background larval competition, (A) social females were similarly likely to oviposit but (B) strongly with-held total number of eggs deposited relative to solitary females. (C) At finer spatial scales, social females like the solitary ones preferentially deposited eggs in pools with larval competitors than without (indicated by OAI values consistently greater than 0). Furthermore, social females were attracted to high larval competitor pools (high positive OAI) despite the availability of a risk-free pool.

#### Finer spatial scale

Neither social status nor level of larval competition influenced the degree of choosiness (OAI) (risk level: *F*-value = 3.026, *p* = 0.08529; status: *F*-value = 0.3696, *p* = 0.54471) or the relationship between OAI and total fecundity (Table 3). On an average, ovipositing females preferred to deposit eggs in pools with larval competitors than without (Fig. 2; mean OAI: 0.1448, bootstrapped CI = 0.169, 0.477). Individuals that chose this strategy laid fewer total number of eggs, than those that distributed equally across pools (Fig. 3, Table 3, glm OAI squared term, *F*=7.0412, df=1, *p <* 0.005).

**Figure 2:**
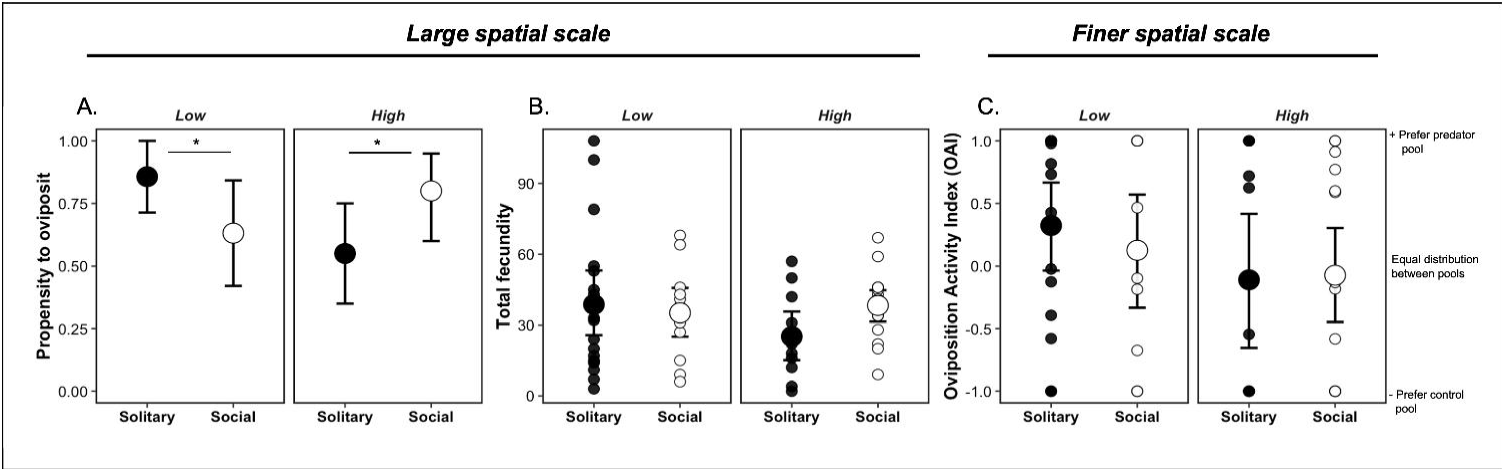
In ‘*Low*’ background predation, (A) social females were less likely to oviposit eggs (B) but laid similar, total number of eggs relative to solitary adults. (C) Here, social females preferred to distribute eggs across pools while solitary females were attracted to low predation pools despite availability of a risk-free pool. In contrast, in ‘*High*’ background predation, (A) social females were more likely to oviposit and (B) laid greater number of eggs relative to solitary adults but (C) distributed their eggs between pools like their solitary counterparts (indicated by OAI values close to 0).

**Figure 3:**
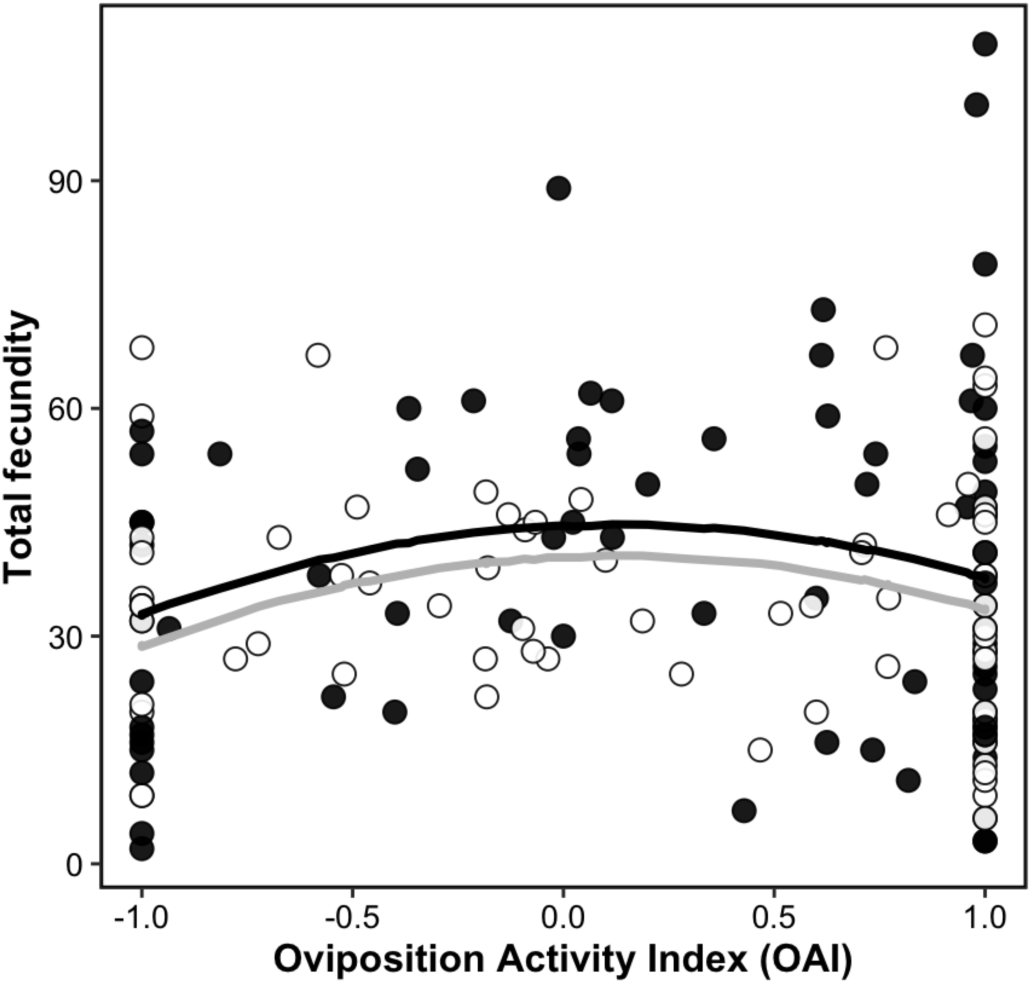
Regardless of social information, and level or type of background risk (predation or competition), females laid fewer eggs in total when they laid all eggs in one pool (control/treatment) but laid greater number of eggs when they distributed eggs between the two pools. Lines represent predictions from the GLMM.

**Table 1:**
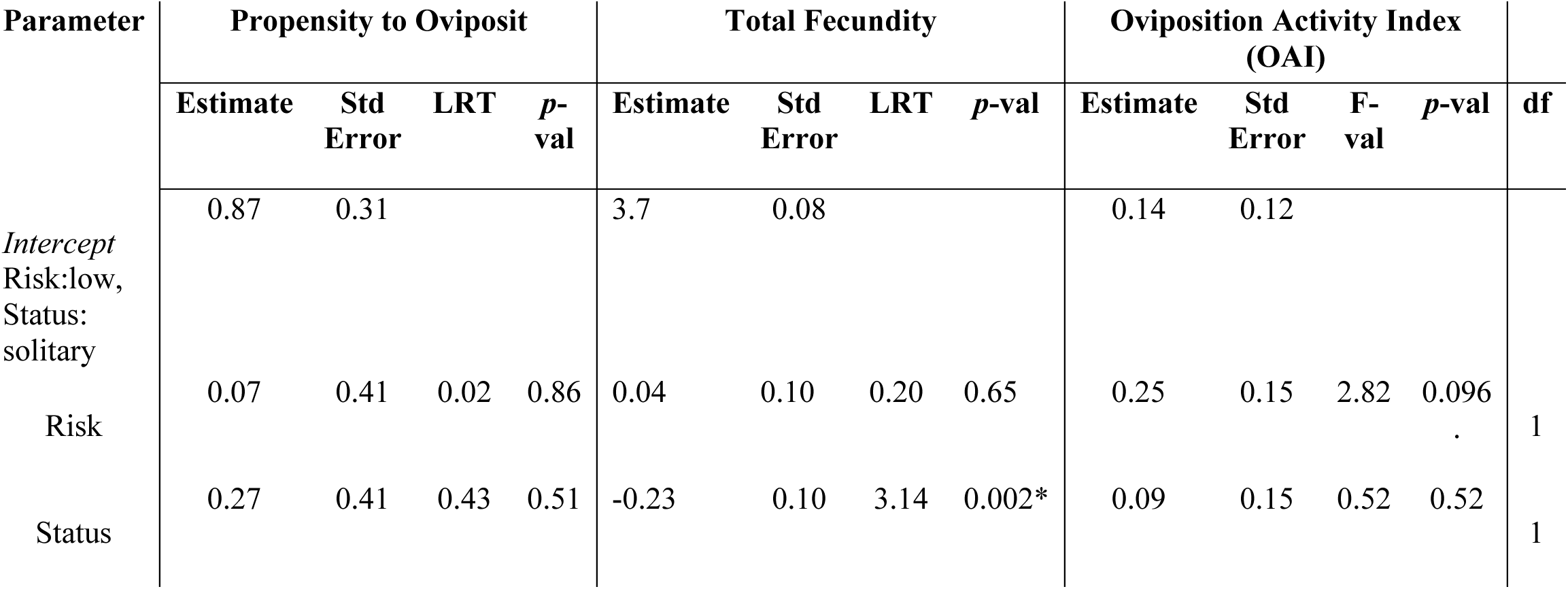
Estimates and statistical significance of variables included in the analyses of (A) Propensity to Oviposit (B) Total Fecundity and (C) Oviposition Activity Index (OAI) of solitary and social individuals (status) in response to varying levels of larval competition (type). Likelihood ratio tests (X^2^) are used to test the significance of fixed effects. Model reduction to obtain the minimal model was performed only if interaction between explanatory variables were non-significant. For variables included in the minimal model, likelihood ratio test statistic from comparison of final model with a model excluding the specified variable is reported.

**Table 2:**
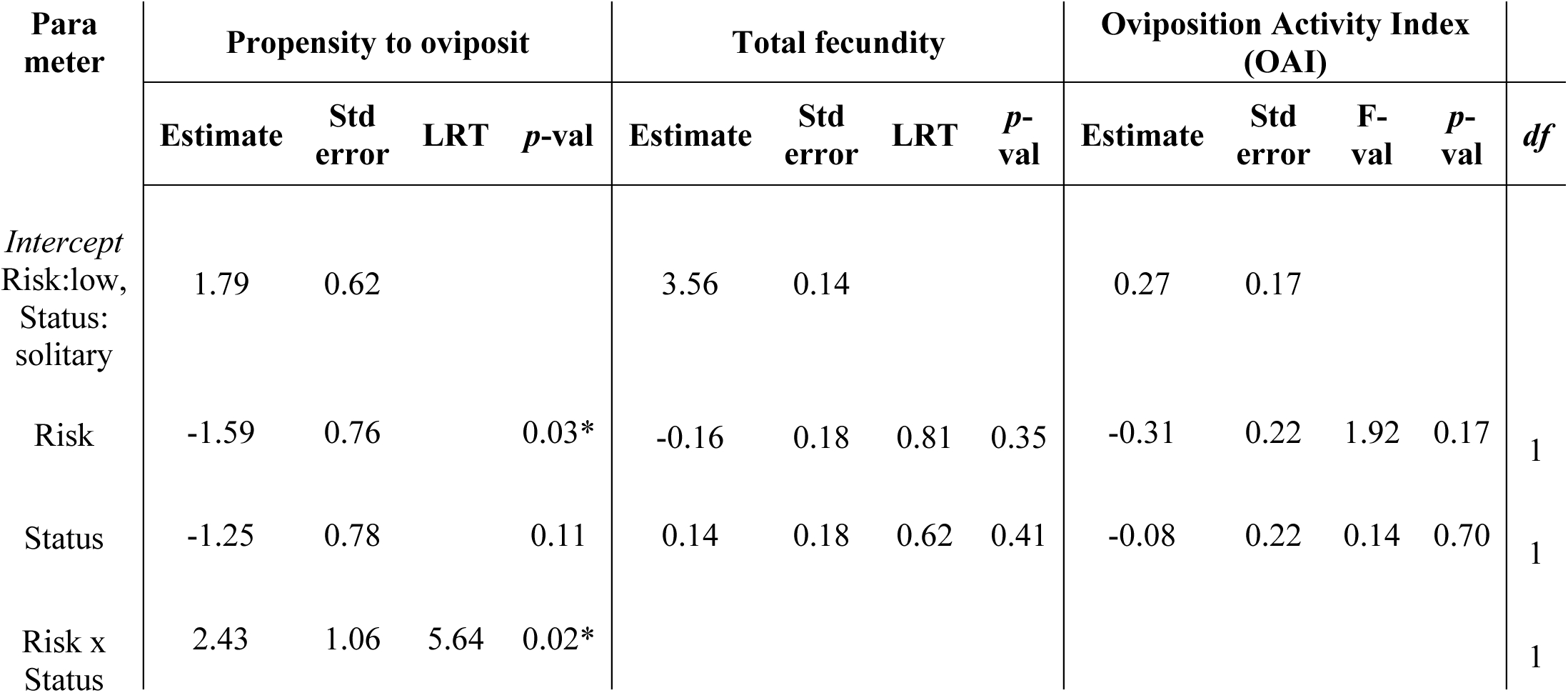
Estimates and statistical significance of variables included in the analyses of (A) Propensity to Oviposit (B) Total Fecundity and (C) Oviposition Activity Index (OAI) of solitary and social individuals (status) in response to varying levels of larval predation (type). Likelihood ratio tests (X^2^) are used to test the significance of fixed effects. Model reduction to obtain the minimal model was performed only if interaction between explanatory variables were non-significant. For variables included in the minimal model, likelihood ratio test statistic from comparison of final model with a model excluding the specified variable is reported.

**Table 3:** Analysis of deviance in the total eggs laid in relation to oviposition activity index (OAI), quadratic term of OAI, social status, and level of risk. Likelihood ratio tests are shown for the terms from a generalized linear model fitted to the total eggs laid.

| Parameter | Coefficient | Std. Error | *df* | *F* | *p* value |
| --- | --- | --- | --- | --- | --- |
| Intercept (Risk: Low,<br>Status: Solitary) | 44.5749 | 3.553 |  |  |  |
| OAI | 2.402 | 2.0477 | 1 | 1.173 | 0.243 |
| OAI^2 | -9.2704 | 3.9718 | 1 | -2.334 | 0.021 * |
| Risk: Social | -4.1033 | 3.1742 | 1 | -1.293 | 0.198 |
| Status: High | -0.1903 | 3.1937 | 1 | -0.060 | 0.953 |

### Larval Predation Risk

#### Larger spatial scale

Overall, both social status and the level of risk significantly influenced the propensity to oviposit but in opposing directions (Fig. 2A, risk level x status: *X^2^* = 5.643, *p* = 0.02178). In ‘*Low*’ background predation, social females were 0.73 times more likely to reject the pool network than solitary females. However, as background larval predation intensified, social females were 1.45 times more likely to accept the pool network. Yet, social information had no detectable effect on the total fecundity of ovipositing females across varying background larval predation (Fig. 2B, risk level: *X^2^* = 0.81506, *p* = 0.352; status: *X^2^* = 0.62340, *p* = 0.416).

#### Finer spatial scale

Neither social status nor level of larval predation had a detectable effect on the degree of choosiness (OAI) (Figure 3, risk level: F-value = 1.9275, p = 0.171; status: F-value = 0.146, p = 0.704) or the relationship between OAI and total fecundity (Table 3). On average, ovipositing females tended to distribute their eggs between predator pools and no predator pools (Figure 4; mean OAI: 0.0875, bootstrapped CI = −0.136, 0.302). Individuals that chose this strategy maximized the total number of eggs laid, compared to those that laid all eggs in one pool (control/ treatment) (Figure 3, Table 3, glm OAI squared term, *F*=7.0412, df=1, *p <* 0.005).

## DISCUSSION

How does social information influence individual egg-laying decisions when oviposition patches pose threat to the progeny? To answer this question, we investigated the oviposition ecology of the mosquito *A aegypti.* We tested how social information (or lack thereof) influences oviposition decisions in ovipositing females across two sources of risk – larval competition and predation. Our findings reveal that social information exerts a stronger influence on egg-laying behavior at broader than finer spatial scales. Social information led to rejecting pool networks under low background predation but prompted acceptance at higher predation rates, contrasting with solitary females (Fig. 1). Additionally, while social cues did not prompt rejection of pool networks amidst background competition, social females deposited fewer eggs, demonstrating social inhibition (Fig. 2). At finer spatial scales, social information modified oviposition behaviour but not always in the direction predicted despite potential negative fitness consequences for its offspring. For instance, females exhibited a preference for ovipositing in pools with conspecific larvae but the presence of larval predation prompted to adopt bet-hedging strategies. Finally, linking the larger to finer spatial scale revealed that regardless of social information females laid fewer eggs in total when they laid all eggs in one pool (control/treatment) but laid greater number of eggs when they distributed eggs between the two pools (Fig. 3). Our study provides evidence that the effect of social information – facilitation, inhibition or negligible effect – both in magnitude and direction is risk and spatial scale dependent. Through our study-design we provide promising evidence for influence of social information alone on intra-individual variation in egg-laying, and potential proximate mechanisms to understand seasonal patterns of oviposition.

### Larval competition

Social information via adult conspecifics strongly modified oviposition behaviour in response to larval competition. Overall, social information had no effect on the propensity to oviposit in a pool network. However, in the presence of conspecifics, individual females strongly withheld more eggs when compared with solitary females. Therefore, females are likely assessing the future competition for their offspring prior to laying eggs. At a finer spatial scale, regardless of social status, females generally preferred to lay eggs in pools with conspecifics. Here, social females showed a slightly stronger preference to oviposit in pools containing high density of larvae as was observed in the field (Edgerly et al 1998). Thus, social information strongly inhibited egg-laying at larger spatial scales but skewed preference to seemingly risky pools at finer scales.

So why do social females strongly prefer to lay eggs in high larval pools given the potential threat to progeny via larvae and eggs of neighbouring competitors? First, the eggs laid may not be in direct competition with larvae currently in water. Thus, females may be accounting for the time lag between larval metamorphosis to optimize egg-laying. Studies on mosquito species - *A triseriatus*, Culex *spp*., and sandfly showed that females show positive oviposition response to habitats with higher egg-densities supporting this explanation (Elnaiem and Ward 1991; Edgerly et al 1998). Focal females here can viewed as the first to enter the patch, and if eggs hatch in cohorts, their progeny would not be in competition with the larvae in water, or eggs laid in the future. Second, high larval density may be an indicator of habitat stability and suitability (Wong et al 2011; Zahiri et al 1997). Further, habitat stability may be strengthened by nutrient availability. Since the pool previously housed a large number of larvae and if nutrient recycling through debris, larval moults etc. results in a nutrient rich pool, treatment pools may signal higher habitat quality (Walker et al 1991). Finally, larval aggregation can prove beneficial by preventing the formation of scum in water thereby reducing predation and parasitism of larvae and reinforcing habitat stability (Pile 1987; Edgerly et al 1998). Thus, in many ways, larval competitors alongside adult conspecifics could be additional sources of social information for ovipositing females.

### Larval predation

Social information strongly modified oviposition behaviour at a large spatial scale in response to larval predation risk. When background predation was low, social information promoted individuals to strongly reject pool networks; and when they do oviposit, to bet-hedge across the control and treatment pools. This is consistent with field patterns measuring oviposition behaviour of *Aedes vexens* with varying background predation risk broadly indicate that females consistently lay eggs in both predator present and predator-free pools across the season (Sharma and Isvaran 2019; Silberbush et al 2014). Interestingly, at a finer spatial scale, ovipositing solitary females preferentially deposited eggs in low predator pools despite the availability of a risk-free pool. When females lacked social information via adult conspecifics, it appears that they made seemingly riskier choices in the laboratory (Silberbush and Blaustein 2011). Empirical evidence from other ovipositing insects demonstrate that although sometimes preferences are suboptimal, they are perhaps better than making random choices (Heard, 1994). The risk to offspring may be mitigated if low predation can rescue offspring from negative-density dependent effects of competition (Albeny-Simoes *et al* 2014). Alternately, if risk-free pools cannot sustain high larval density, then asymmetric egg-laying skewed to low predator pools may result in higher fitness.

In contrast, when background predation was high, social information strongly promoted individuals to oviposit in pool networks. However like earlier, on average, at finer spatial scales, social females tended to spread eggs between no predator and high predator pools. Individuals choosing this strategy laid larger numbers of eggs in the pool-network, compared to those that laid all their eggs in one pool. In line with our hypothesis, oviposition in high predator pool can possibly be explained if presence of additional unrelated larvae contributed by other females in the environment results in a potential dilution effect, thus reducing per capita risk. (Albeny-Simoes *et al* 2014). Further studies using field mesocosms could validate if egg-laying at the population scale is skewed towards high predator pools. Together, these findings highlight how social information shapes adaptive egg-laying behaviors in response to predation threats.

### Social clues in risky pools

By tracking individual oviposition behaviour in a group setting using a rare experimental approach, we provide unique insight about the behavioral ecology and sociobiology of asocial insects like mosquitoes. First, our study shows that social information via presence of neighbouring conspecifics alone can result in individual females exhibiting preference to riskier patches (Albeny-Simoes *et al* 2014). When background competition and predation risk was high, social information facilitated oviposition behaviour albeit at different spatial scales. In high predation, social information facilitated egg-laying at scale of pool-network but females bet-hedged across predator and no predator pools; while in high competition, it facilitated egg-laying in seemingly risky high competition pools over no risk at fine scales (*reviewed in* Buxton et al 2020). Although, social information did not operate in the same expected direction across both scales, it provides evidence that social information can facilitate oviposition behaviour in the background of risky patches (Prokopy and Duan 1998). Second, we demonstrate the influence of social cues on intra-individual variation in egg laying using the unique mosquito biology. This rare approach allows us to capture the oviposition response of individuals in a group setting, which is often undisclosed due to limitations in study-design in other model systems (but *see* Kelly et al 2018). We find that individuals even in a social context, can completely avoid ovipositing in the pool-network despite availability of risk-free options. Thus, individuals choose not to make a choice. Of the females that do choose to oviposit, individuals exhibit a wide range of behaviors – bet-hedging to laying all her eggs in the risky, or risk-free pool just like her solitary counterparts. We speculate that uncertainty in the behaviour decision of other mosquitoes (i.e. future competition), or uncertainty in the cost to benefit ratio of offspring performance across the pools may elicit a wide range of responses from an individual (Albeny-Simoes *et al* 2014). Our study allows to capture the suite of behaviors that individuals in a group can exhibit despite potentially staggering fitness differences incurred by their offspring. Finally, individuals in a group rely on integrating public and personal information to make oviposition decisions (Malek and Long 2020). *A. aegypti* females procure personal information about the pool quality by using chemical receptors at the back of their legs to skim the water surface. If individuals relied only on personal information, then individuals in a group would behave identical to solitary individuals in our experiment. Similarly, if individuals relied only on public information i.e. presence of adult conspecifics, then all individuals in a social setting regardless of risk type should behave similarly. Our study does not provide support for either hypothesis. Rather, we observe the distinct behavioral choices across both type and level of risk when individuals are in a social setting. In the wild, mosquitoes may be required to assess the type of risk in a patch and calibrate the anticipated egg-laying based on neighbouring adult conspecifics prior to making choices. Thus, decision making in mosquitoes maybe more complex than previously anticipated.

Four other aspects of behavioural ecology and sociobiology of mosquitoes could enhance our understanding of the cause and consequences of their oviposition behaviour in complex landscapes. First, future experiments may benefit from comparing oviposition of solitary breeding, solitary breeding with perceived conspecific risk, and “grouped” all breeding females. Our results may reflect the behaviour of the first ovipositing female in a group of females as seen in certain parasitoid species (Godfray et al 1994). Egg-laying choices by subsequent females or a group can offer more comprehensive insights into the effects of social information at a fine spatial scale. Thus, the emergent behaviour of ovipositing females in a social setting can be informative of the consequences at finer spatial scales. Second, parameterizing the effects of increased conspecific densities on oviposition choices can map behaviour mechanisms to outcomes of temporal egg-densities. For the current experiment we chose densities reflective of minimum (1 mosquito) and maximum (4 mosquitoes) on average per ovitrap recorded via capture techniques on field sites across seasons (Lourenço-de-Oliveira 2008). Third, corroborating lab experimental data with manipulated field mesocosms can provide robust evidence of influence of social cues along with other environmental variables like pool-size and desiccation risk on spatial aggregation and fine scale choices of mosquitoes. Finally, experiments to measure fitness consequences of these outcomes can provide insight into cost to benefit ratio that can help explain anomalous outcomes at finer spatial scales. Furthermore, given the importance of *A. aegypti* as disease vectors, future studies could explore how their trait responses also influence choices, disease dynamics and life history traits (Ramesh & Hall, 2023).

Together, our findings reveal that social information exerts a stronger influence on egg-laying behavior at larger spatial scales, *i.e*., at the scale of pool networks, than between neighbouring pools. However, the direction of that modification was context dependent and not often in the direction predicted despite potential negative fitness consequences for its offspring. with increasing larval predation facilitating oviposition but larval competition imposing inhibition. Then, our study contextualizes key findings from the past two decades of OSS experimentation, suggesting that social facilitation, inhibition, or no effect on oviposition behavior may arise primarily due to perceived conspecific rivals, varying with offspring risk and spatial scale. Mechanistic models can, in the future, integrate individual behavioural changes influenced by social information. By linking socio-oviposition behavior to population dynamics we can better predict seasonal spread of mosquitos and highlight the implications for vector transmission.

## Acknowledgements

AR was supported by the Scholarship for Higher Education (SHE) INSPIRE Fellowship at IISER Pune, Department of Science and Technology (DST), DBT-IISc Partnership Programme and a MSU Presidential Postdoctoral Fellowship during the completion of this manuscript. We thank D Barua for helpful discussion on the manuscript. AR credits Y Chryssomallis and C Raj for artistic inspiration.

## REFERENCES

Abernathy, R. L., Teal, P. E., & Tumlinson, J. H. (1994). Age and crowding affects the amount of sex pheromone and the oviposition rates of virgin and mated females of Helicoverpa zea (Lepidoptera: Noctuidae). Annals of the Entomological Society of America, 87(3), 350–354.

Aikins, C., Altizer, S., & Sasaki, T. (2023). Neither copy nor avoid: no evidence for social cue use in monarch butterfly oviposition site selection. Journal of Insect Behavior, 36(1), 33–44.

Bates, D., Maechler, M., Bolker, B., Walker, S., Christensen, R. H. B., Singmann, H., … & Bolker, M. B. (2015). Package ‘lme4’. convergence, 12(1), 2.

Betts, M. G., Hadley, A. S., Rodenhouse, N., & Nocera, J. J. (2008). Social information trumps vegetation structure in breeding-site selection by a migrant songbird. Proceedings of the Royal Society B: Biological Sciences, 275(1648), 2257–2263.

Blaustein, L., Kiflawi, M., Eitam, A., Mangel, M., & Cohen, J. E. (2004). Oviposition habitat selection in response to risk of predation in temporary pools: mode of detection and consistency across experimental venue. Oecologia, 138, 300–305.

Bonnie, K. E., & Earley, R. L. (2007). Expanding the scope for social information use. Animal Behaviour, 74(2), 171–181.

Buxton, V. L., Enos, J. K., Sperry, J. H., & Ward, M. P. (2020). A review of conspecific attraction for habitat selection across taxa. Ecology and evolution, 10(23), 12690–12699.

Chess, K. F., Ringo, J. M., & Dowse, H. B. (1990). Oviposition by two species of Drosophila (Diptera: Drosophilidae): behavioral responses to resource distribution and competition. Annals of the Entomological Society of America, 83(4), 717–724.

Clements, A. N. (1992). Development, nutrition and reproduction. The biology of mosquitoes, 1, 509.

Drugowitsch, J., Moreno-Bote, R., Churchland, A. K., Shadlen, M. N., & Pouget, A. (2012). The cost of accumulating evidence in perceptual decision making. Journal of Neuroscience, 32(11), 3612–3628.

Edgerly, J. S., McFarland, M., Morgan, P., & Livdahl, T. (1998). A seasonal shift in egg-laying behaviour in response to cues of future competition in a treehole mosquito. Journal of Animal Ecology, 67(5), 805–818.

Elnaiem, D. E. A., & Ward, R. D. (1991). Response of the sandfly Lutzomyia longipalpis to an oviposition pheromone associated with conspecific eggs. Medical and veterinary entomology, 5(1), 87–91.

Gibbs, M., Lace, L. A., Jones, M. J., & Moore, A. J. (2005). Egg size-number trade-off and a decline in oviposition site choice quality: female Pararge aegeria butterflies pay a cost of having males present at oviposition. Journal of Insect Science, 5(1), 39.

Gimnig, J. E., Ombok, M., Otieno, S., Kaufman, M. G., Vulule, J. M., & Walker, E. D. (2002). Density-dependent development of Anopheles gambiae (Diptera: Culicidae) larvae in artificial habitats. Journal of Medical Entomology, 39(1), 162–172.

Godfray, H. C. J. (1994). Chapter 3: Oviposition in Parasitoids: behavioral and evolutionary ecology. Princeton University Press.

Heard, S. B. (1994). Imperfect oviposition decisions by the pitcher plant mosquito (Wyeomyia smithii). Evolutionary Ecology, 8, 493–502.

Kelly, J. K., Chiavacci, S. J., Benson, T. J., & Ward, M. P. (2018). Who is in the neighborhood? Conspecific and heterospecific responses to perceived density for breeding habitat selection. Ethology, 124(4), 269–278.

Le Goff, G. J., Mailleux, A. C., Detrain, C., Deneubourg, J. L., Clotuche, G., & Hance, T. (2010). Group effect on fertility, survival and silk production in the web spinner Tetranychus urticae (Acari: Tetranychidae) during colony foundation. Behaviour, 147(9), 1169–1184.

Lehmann, T., Dao, A., Yaro, A. S., Diallo, M., Timbiné, S., Huestis, D. L., … & Traoré, A. I. (2014). Seasonal variation in spatial distributions of Anopheles gambiae in a Sahelian village: evidence for aestivation. Journal of medical entomology, 51(1), 27–38.

Lourenço-de-Oliveira, R., Lima, J. B. P., Peres, R., da Costa Alves, F., Eiras, Á. E., & Codeço, C. T. (2008). Comparison of different uses of adult traps and ovitraps for assessing dengue vector infestation in endemic areas. Journal of the American Mosquito Control Association, 24(3), 387–392.

Malek, H. L., & Long, T. A. (2020). On the use of private versus social information in oviposition site choice decisions by Drosophila melanogaster females. Behavioral Ecology, 31(3), 739–749.

Miller, C. W., Fletcher Jr, R. J., & Gillespie, S. R. (2013). Conspecific and heterospecific cues override resource quality to influence offspring production. PLoS One, 8(7), e70268.

Munga, S., Minakawa, N., Zhou, G., Barrack, O. O. J., Githeko, A. K., & Yan, G. (2014). Effects of larval competitors and predators on oviposition site selection of Anopheles gambiae sensu stricto. Journal of medical entomology, 43(2), 221–224.

Mukherjee, S., & Blaustein, L. (2019). Effects of predator type and alternative prey on mosquito egg raft predation and destruction. Hydrobiologia, 846(1), 215–221.

Pile, M. M. (1987). Pheromone-mediated Behaviour of the Mosquito “Culex Quinquefasciatus”: Elicited by an Oviposition Attractant and Its Derivatives (Doctoral dissertation).

Prokopy, R. J., & Bush, G. L. (1973). Oviposition by grouped and isolated apple maggot flies. Annals of the Entomological Society of America, 66(6), 1197–1200.

Prokopy, R. J., & Duan, J. J. (1998). Socially facilitated egglaying behavior in Mediterranean fruit flies. Behavioral ecology and sociobiology, 42, 117–122.

Ramesh, A., & Hall, S. R. (2023). Niche theory for within-host parasite dynamics: Analogies to food web modules via feedback loops. Ecology Letters, 26(3), 351–368.

Refsnider, J. M., & Janzen, F. J. (2010). Putting eggs in one basket: ecological and evolutionary hypotheses for variation in oviposition-site choice. Annual Review of Ecology, Evolution, and Systematics, 41, 39–57.

Resetarits Jr, W. J., & Wilbur, H. M. (1989). Choice of oviposition site by Hyla chrysoscelis: role of predators and competitors. Ecology, 70(1), 220–228.

Sharma, M., & Isvaran, K. (2019). Trait evolution under multiple selection pressures: Prey responses to predictable and unpredictable variation. bioRxiv, 816314.

Sharma, M., Quader, S., Guttal, V., & Isvaran, K. (2020). The enemy of my enemy: multiple interacting selection pressures lead to unexpected anti-predator responses. Oecologia, 192, 1–12.

Silberbush, A., & Blaustein, L. (2011). Mosquito females quantify risk of predation to their progeny when selecting an oviposition site. Functional Ecology, 25(5), 1091–1095.

Silberbush, A., Tsurim, I., Margalith, Y., & Blaustein, L. (2014). Interactive effects of salinity and a predator on mosquito oviposition and larval performance. Oecologia, 175, 565–575.

Team, R. C. (2021). R: A language and environment for statistical computing. R Foundation for Statistical Computing.

Tóth, Z., Jaloveczki, B., & Tarján, G. (2020). Diffusion of social information in non-grouping animals. Frontiers in Ecology and Evolution, 8, 586058.

Visser, M. E. (1995). The effect of competition on oviposition decisions of Leptopilina heterotoma (Hymenoptera: Eucoilidae). Animal Behaviour, 49(6), 1677–1687.

Walker, E. D., Lawson, D. L., Merritt, R. W., Morgan, W. T., & Klug, M. J. (1991). Nutrient dynamics, bacterial populations, and mosquito productivity in tree hole ecosystems and microcosms. Ecology, 72(5), 1529–1546.

Wong, J., Stoddard, S. T., Astete, H., Morrison, A. C., & Scott, T. W. (2011). Oviposition site selection by the dengue vector Aedes aegypti and its implications for dengue control. PLoS neglected tropical diseases, 5(4), e1015.

Zahiri, N., Rau, M. E., & Lewis, D. J. (1997). Oviposition responses of Aedes aegypti and Ae. atropalpus (Diptera: Culicidae) females to waters from conspecific and heterospecific normal larvae and from larvae infected with Plagiorchis elegans (Trematoda: Plagiorchiidae). Journal of medical entomology, 34(5), 565–568.

